# Glutamatergic neuron types in the amygdala of the urodele amphibian Pleurodeles waltl

**DOI:** 10.1101/2022.06.15.496313

**Authors:** Astrid Deryckere, Jamie Woych, Eliza C. B. Jaeger, Maria Antonietta Tosches

**Affiliations:** Department of Biological Sciences, Columbia University; New York City, 10027 New York, USA

**Keywords:** Amygdala, Amphibian, Neuron types, Brain Evolution

## Abstract

The amygdala is a complex brain structure in the vertebrate telencephalon, essential for regulating social behaviors, emotions and (social) cognition. In contrast to the vast majority of neuron types described in the many nuclei of the mammalian amygdala, little is known about the neuronal diversity in non-mammals, making reconstruction of its evolution particularly difficult. Here, we characterize glutamatergic neuron types in the amygdala of the salamander *Pleurodeles waltl*. Our single-cell RNA sequencing data indicate the existence of at least ten distinct types and subtypes of glutamatergic neurons in the salamander amygdala. *In situ* hybridization for marker genes indicates that these neuron types are located in three major subdivisions: the lateral amygdala, the medial amygdala, and a newly-defined area demarcated by high expression of the transcription factor *Sim1*. The gene expression profiles of these neuron types suggest similarities with specific neuron types in the sauropsid and mammalian amygdala, and in particular the evolutionary conservation of *Sim1*-expressing amygdalar neurons in tetrapods. Taken together, our results reveal a surprising diversity of glutamatergic neuron types in the amygdala of salamanders, despite the anatomical simplicity of their brain.

## Introduction

The amygdala is a complex brain region that controls a variety of behaviors crucial for survival (Swanson and Petrovich 1998). In mammals, this part of the brain not only controls fear responses (i.e. escape) through direct connections with motor and autonomic systems, but it is also critical for social behaviors and associative learning. The amygdala develops in the caudal telencephalon, and includes nuclei derived from pallium and subpallium (Aerts and Seuntjens 2021). Furthermore, the amygdala harbors cells that migrate in from the hypothalamus and the prethalamic eminence (García-Moreno et al. 2010; Garcia-Calero et al. 2021; Morales et al. 2021). As a result of their complex developmental history, amygdalar nuclei have distinct neurochemical, hodological, and functional properties. Anatomical and molecular studies in mice suggest a great diversity of amygdalar neuron types; however, the characterization of cellular heterogeneity remains limited to a few nuclei and to the sampling of small numbers of cells (Wu et al. 2017; Zeisel et al. 2018; O’Leary et al. 2020).

Telencephalic areas with hodological and molecular similarities to the mammalian amygdala have been clearly identified in representatives of all the main classes of jawed vertebrates (Medina et al. 2017) (data on agnatha remain scarce). Therefore, the amygdala as a whole seems to be ancient in vertebrates, in line with its crucial functions. However, several questions on the evolution of the amygdala remain open. What are the exact boundaries and subdivisions of the amygdala in non mammals? How do amygdalar subdivisions relate to each other across species? And ultimately, how did the mammalian amygdala, with its complex organization, emerge?

Unsurprisingly, the most controversial points concern the pallial amygdala, because the evolution of the pallium as a whole remains unclear. The well-known tetrapartite model for the pallium, introduced by Puelles and colleagues, postulated that the pallial amygdala is part of the so-called ventral pallium, the pallial region defined by the absence of expression of the transcription factor *Emx1* in its ventricular zone (Fernandez et al. 1998; Puelles et al. 2000; Gorski et al. 2002; Brox et al. 2004). In the more recent versions of this model, the mammalian ventral pallium includes the pallial amygdala, the piriform cortex, and the ventral endopiriform nucleus, among others (Laberge et al. 2006; Puelles 2017). Following this model, a ventral pallium has been identified in other vertebrates on the basis of topological criteria and conserved developmental gene expression patterns. In tetrapods, it is clear that the ventral pallium includes amygdalar regions, in addition to other neural territories with controversial homologies. For example, the reptilian ventral pallium is heterogeneous: it includes the lateral cortex (an olfactory-recipient cortical region, considered the homolog of the mammalian piriform cortex) and the dorsal ventricular ridge (DVR), comprising anterior and posterior subdivisions (aDVR and pDVR). In birds, the nidopallium and arcopallium would correspond to the aDVR and pDVR, respectively (Medina et al. 2021). While the homology of the pDVR and of the arcopallium with parts of the mammalian pallial amygdala has been reconciled, the nature of the aDVR and of the nidopallium remains debated (Bruce and Neary 1995; Briscoe and Ragsdale 2018; Colquitt et al. 2021; Gedman et al. 2021; Medina et al. 2021; Tosches 2021).

Acknowledging the important differences in gene expression profiles in the ventral pallium, Medina and colleagues proposed the partition of the original ventral pallium in two pallial sectors; the ventral and ventrocaudal pallium. The latter sector includes the pallial amygdala (Medina et al. 2017; Desfilis et al. 2018). Taking these observations a step further, the new radial model for the pallium proposed by Puelles and colleagues abandons the classical tetrapartite scheme, and posits that the mammalian pallial amygdala arises from radial segments that are completely distinct from the rest of the pallium, which now takes the name of cortical pallium (Puelles et al. 2019).

As the sister group of amniotes, amphibians are extremely informative to reconstruct brain evolution in tetrapods. The amphibian pallium has a simple organization, as it consists of a periventricular cell layer and a plexiform layer, and is devoid of clearly distinct nuclei. Subdivisions of the amphibian pallium have been identified with the combination of tracing, neuroanatomical, and molecular studies. In amphibians, the terms medial, dorsal, lateral, and ventral pallium have been used to label these pallial subdivisions, and in this article, we will use this traditional terminology (González et al. 2020). However, it is important to highlight that these terms have not been updated to follow the changing tetrapartite model for the pallium. For example, new data suggests the homology of the urodele lateral pallium with bowl cells in the reptilian lateral cortex and with semilunar cells in the mammalian piriform cortex; therefore in amphibians the term “lateral pallium” refers to neurons homologous to neurons in the amniote ventral pallium, and not in the amniote lateral pallium (Wullimann 2017; Tosches 2021; Lust et al. 2022; Woych et al. 2022).

The neuroanatomical heterogeneity of the amphibian ventral pallium was already noticed by several investigators, who distinguished subdivisions of this area along the medio-lateral axis even before the term ventral pallium was introduced (Northcutt and Kicliter 1980; Neary 1990; Bruce and Neary 1995). For example, Neary identified three regions of the “lateral pallium”, corresponding to the lateral and ventral pallium in modern terms (Neary 1990), and in addition Marín described a “subpallial-pallial transition area” (SPTA) (Marín et al. 1997b, 1997a). Molecular data confirmed the existence of a “ventral subdivision of the ventral pallium” of an amygdalar nature; this region was named “lateral amygdala” by Moreno, Gonzales and colleagues (Moreno and González 2004; Moreno et al. 2004). The lateral amygdala is one of the three amygdalar subdivisions identified so far in amphibians; the other two, of subpallial origin, are the medial and the central amygdala (Laberge et al. 2006; Hall et al. 2013; González et al. 2020). In *Xenopus*, the lateral amygdala is demarcated by the expression of the transcription factor *Lhx9*, and at mid-telencephalic levels it is nested between the rest of the ventral pallium dorsally and the striatum ventrally (Brox et al. 2003; Moreno et al. 2004). At caudal levels, the organization of the amphibian amygdala remains less clear. In urodeles, Northcutt and Kicliter (Northcutt and Kicliter 1980) identified an “amygdala pars lateralis” and proposed that this area has a pallial origin. However, Moreno and Gonzales (Moreno and González 2007a) found expression of subpallial markers in the caudal telencephalon, and renamed this region medial amygdala, proposing its homology with the mammalian medial amygdala.

These observations raise several questions about the organization of the amygdala in amphibians, and the evolution of the amygdala in tetrapods. First, is the new radial model of the pallium applicable to amphibians? The model proposes the existence of distinct radial sectors for the pallial amygdala and the cortical pallium. To confirm the existence of one (or multiple) pallial amygdala radial sector(s) in amphibians, it is necessary to clarify the boundaries with the ventral pallium and with the subpallium along the entire rostro-caudal axis of the telencephalon. Second, is the amphibian pallial amygdala heterogeneous, as it is in sauropsids and mammals? Recent work by Garcia-Calero, Puelles and colleagues (Garcia-Calero et al. 2020) demonstrated that the mammalian pallial amygdala develops from five radial sectors, which give rise to amygdalar nuclei with distinct gene expression profiles and connectivity. A single-cell RNA sequencing (scRNAseq) analysis of the turtle pallium identified five distinct clusters of single cell transcriptomes (a proxy for cell types) in the pDVR, which were mapped to distinct subregions of the pDVR using *in situ* hybridization for marker genes (Tosches et al. 2018). In contrast, it is unclear whether the amphibian pallial amygdala includes multiple types of neurons or distinct radial subdivisions.

Finally, does the amphibian amygdala include neurons with extratelencephalic origins, as it is the case in sauropsids and mammals? (García-Moreno et al. 2010; Garcia-Calero et al. 2021; Morales et al. 2021; Metwalli et al. 2022)

To address these questions, here we characterize glutamatergic neuron types in the amygdala of the urodele amphibian *Pleurodeles waltl*. In a recent study, we used scRNAseq to profile the entire telencephalon of *Pleurodeles* (*Woych et al*. *2022*). Building on this resource, we select molecular markers for clusters of amygdalar glutamatergic neurons, and characterize the expression of these markers with *in situ* hybridization. Our results clarify the boundaries of pallial and subpallial amygdala, reveal the existence of cellular heterogeneity in the pallial amygdala, and suggest the presence of a hypothalamus-amygdala migratory stream in amphibians.

## Materials and Methods

### Animals

Adult *Pleurodeles waltl* were obtained from a breeding colony established at Columbia University, and maintained in an aquatics facility at 20°C under a 12L:12D cycle (Joven et al. 2015). All experiments were conducted in accordance with the NIH guidelines and with the approval of the Columbia University Institutional Animal Care and Use Committee (IACUC protocol AC-AABF2564). Experiments were performed with adult (5-19 months) male and female salamanders.

### Single-cell RNA sequencing data analysis

The dataset was generated from single-cell RNA sequencing of the *Pleurodeles waltl* brain as described in (Woych et al. 2022). For in-depth analysis of the amygdala, the data were subsetted, and heterogeneous clusters (TEGLU25, 35 and 38) were subclustered based on differential gene expression. The cluster annotation was updated to reflect the findings from the work described in this paper.

### Tissue preparation

Tissue preparation, *in situ* hybridization, immunochemistry, and imaging of brain sections were performed as described in (Woych et al. 2022) and summarized in brief below. Adult animals were anesthetized by submersion in 0.2% MS-222, perfused with PBS and 4% PFA, and decapitated. Afterwards, brains were extracted, postfixed overnight at 4 °C and washed in PBS.

### Immunohistochemistry

Following tissue preparation, coronal 70 um sections were made with a Leica VT1200S vibratome. The sections were blocked in blocking solution for 1 hr at RT, then incubated in rabbit anti-Foxg1 (1:1000, Abcam ab18259) primary antibody solution for 72 hrs at 4 °C. Tissue was washed and incubated for 2 hr at room temperature in goat anti-rabbit IgG conjugated to Alexa 594 (1:500, Invitrogen). Sections were again washed, counterstained with DAPI, and mounted on glass slides with DAKO fluorescent mounting medium (Agilent Technologies). Imaging was performed at 20x with a confocal microscope (Zeiss LSM800), and images were processed in Fiji.

### Colorimetric *in situ* hybridization

Fragments for ISH were either generated by PCR amplification of ∼1kb sequences from a *Pleurodeles waltl* brain cDNA library, or ordered from Twist Bioscience and cloned into the pCRII-TOPO vector (Invitrogen) with the Hifi DNA Assembly Kit (NEB). Plasmids were verified by Sanger sequencing, and linearized. Anti-sense RNA probes were then generated by *in vitro* transcription using Sp6 polymerase and purified with the Monarch RNA Cleanup Kit (NEB). Primers and gene block sequences are provided in Supplementary Table 1 and 2. Following tissue preparation, the sections were postfixed, and the pia mater removed. The sections were permeabilized with Proteinase K, and acetylated. At 55-62 °C, the sections were pre-hybridized for 1 hr in Hybridization Mix. The sections were then incubated 1-2 nights in Hybridization Mix with denatured riboprobes (1-2 ng/uL). Two low stringency and two high stringency washes at 55-62 °C, and two MABT washes at RT were followed by blocking, and overnight incubation in anti-DIG-Alkaline phosphatase antibody (Roche) solution at 4 °C. Sections were washed, then colorimetric (NBT/BCIP) signal was developed for 1-5 days, with fresh staining solution provided twice a day during development. Staining was stopped by washing in PBS (pH 7.4), and sections mounted on glass slides with DAKO mounting medium. Imaging was performed at 10x with an upright brightfield microscope (Leica DMR with Basler color camera, ACCU-Slide MS software). Image background subtracted using Photoshop PS6.

### Whole mount HCR ISH with iDISCO clearing

Whole brain tissue was stained and cleared as described in (Woych et al. 2022) using a method of iDISCO modified for HCR-v3.0 (Choi et al. 2018). Probe pairs were either ordered from Molecular Instruments or designed using the insitu_probe_generator (Kuehn et al. 2021) and ordered from IDT (probe sequences used for generation of probe sets provided in Supplementary Table 3). Brains were incubated in 2-4 pmol of each probe set (*Slc17a6, Gad1, Sox6, Sim1, Otp*). Imaging was performed using a LaVision Ultramicroscope II light sheet microscope at 4X magnification and 2 μm resolution. Images were visualized using ImarisViewer 9.8.0, and virtually downsized (0.4x) and resliced in Fiji (Schindelin et al. 2012). Sample impurities within the ventricle or on the brain surface were masked and filtered out using ImarisViewer 9.8.0, according to fluorescence intensity.

## Results

### The ventral pallium and the pallial amygdala are molecularly and anatomically distinct in *Pleurodeles waltl*

In a recent scRNAseq study (Woych et al. 2022), we identified 114 clusters of neurons sampled from the adult telencephalon of the urodele amphibian *Pleurodeles waltl*. Of these, 41 clusters were annotated as telencephalic glutamatergic neurons, on the basis of the expression of marker genes including the transcription factors *Foxg1, Emx1, Tbr1*, and *Neurod2*, the glutamate transporters *Slc17a6* and *Slc17a7*, and for the lack of expression of diencephalic markers (Woych et al. 2022). After staining brain sections or whole-mount brain preparations for cluster-specific marker genes, we assigned a regional identity to each of these clusters. We identified clusters that belong to the cortical pallium (TEGLU1-24, Fig. 1A, (Woych et al. 2022)), and here we describe the telencephalic glutamatergic clusters (TEGLU25 and TEGLU33-38) that map to distinct glutamatergic cell types in the *Pleurodeles* amygdala.

**Fig. 1.**
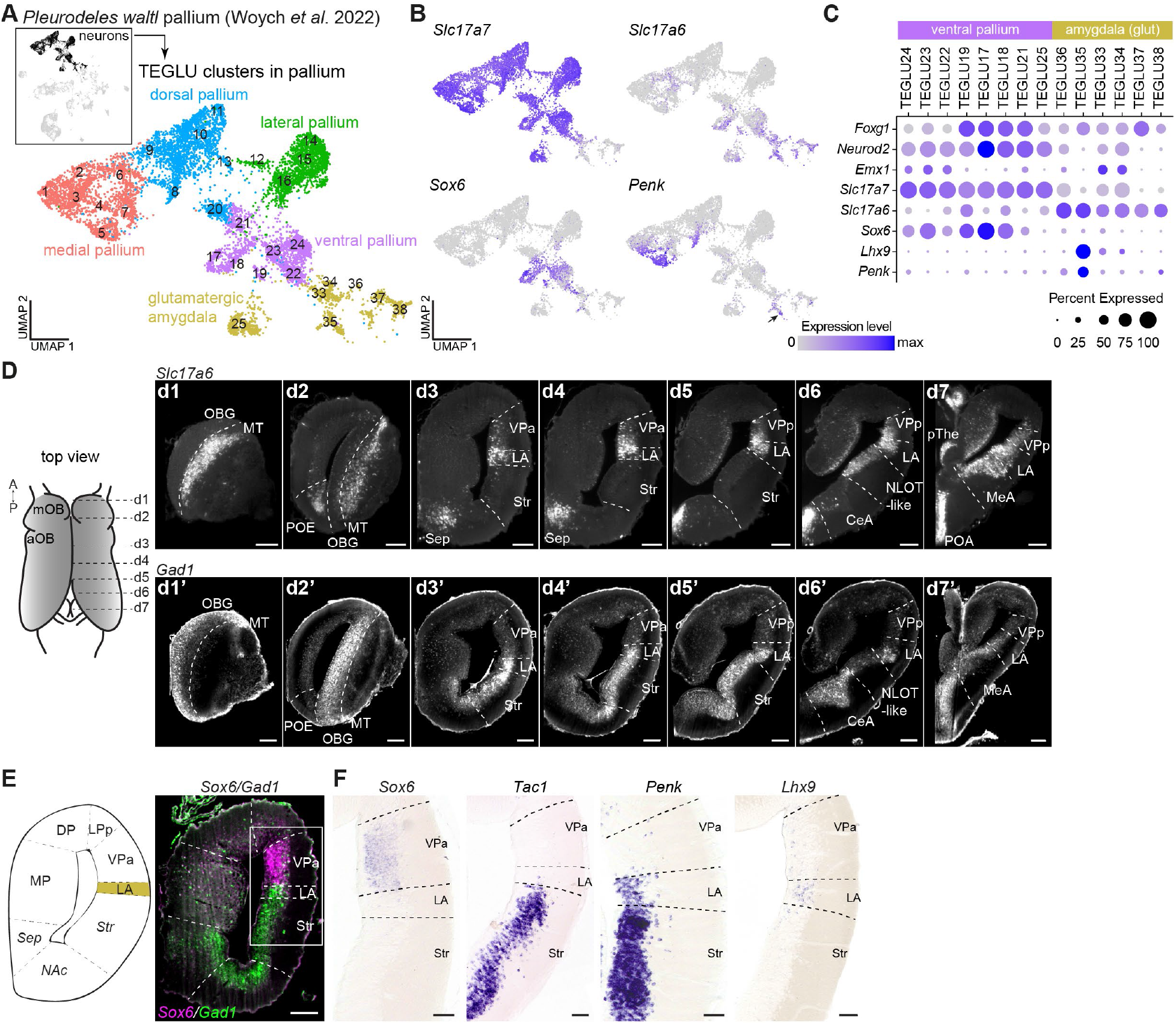
Boundaries of the *Pleurodeles* amygdala territories. **(A)** UMAP plot of 10,125 single-cell transcriptomes of telencephalic glutamatergic neurons, derived from the neuronal dataset presented in Woych et al. 2022 (highlighted in the boxed area). Clusters are annotated by the inferred pallial regions. **(B)** UMAP plots showing single cells colored according to the expression of *Slc17a7* (cortical pallium and medial amygdala), *Slc17a6* (glutamatergic amygdala and parts of the ventral and medial pallium), *Sox6* (ventral pallium) and *Penk* (parts of the glutamatergic amygdala and medial and dorsal pallium). **(C)** Dot plot showing the expression of key marker genes defining ventral pallium and glutamatergic amygdala. **(D)** Left: schematic of the top view of the salamander telencephalon (gray) with dashed lines indicating the optical sectioning planes for images on the right. Right: optical coronal sections after whole-mount HCR ISH, clearing and light sheet imaging for *Slc17a6* (top) and *Gad1* (bottom). Scale bars represent 200 um. **(E)** Left: schematic of a coronal section at mid-telencephalic level with the lateral amygdala territory indicated in yellow. Right: expression of *Sox6* and *Gad1* showing the boundary between the VPa and LA with box indicating the magnified region in F. Scale bar represents 200 um. **(F)** Left to right: expression of *Sox6, Tac1, Penk* and *Lhx9* indicating the borders of the lateral amygdala with the ventral pallium dorsally and striatum ventrally. Scale bars represent 100 um. Abbreviations: aOB, accessory olfactory bulb; CeA, central amygdala; DP, dorsal pallium; LA, lateral amygdala; LPp, posterior lateral pallium; MeA, medial amygdala; mOB, main olfactory bulb; MP, medial pallium; MT, mitral/tufted cells; NAc, nucleus accumbens; NLOT-like, nucleus of the olfactory tract-like; OBG, olfactory bulb GABAergic; POA, preoptic area; POE, postolfactory eminence; pThe, prethalamic eminence; Sep, septum; Str, striatum; TEGLU, telencephalic glutamatergic; VPa, anterior ventral pallium; VPp, posterior ventral pallium.

Glutamatergic neuron types in the cortical pallium and in the amygdala can be distinguished by the expression of genes in different combinations and levels, with only a few individual genes marking either group of neurons. Notably, the vesicular glutamate transporters *Slc17a6* (*Vglut2*) and *Slc17a7* (*Vglut1*) are differentially expressed among telencephalic glutamatergic neurons. Specifically, *Slc17a7* is expressed at high levels in the entire cortical pallium, but also in the amygdala cluster TEGLU25. Clusters TEGLU33 to TEGLU38 do not express *Slc17a7*, but instead express *Slc17a6* at high levels. *Slc17a6* is also found in a few other glutamatergic clusters in the cortical pallium (Fig. 1B-C).

To determine the borders of the amygdala, we characterized the expression of *Slc17a6* and *Gad1* in the telencephalon in three dimensions. We combined whole-brain hybridization chain reaction (HCR) *in situs*, brain clearing with iDISCO, and light-sheet imaging. Here, we present these three-dimensional datasets by slicing them virtually at different angles (Fig. 1D).

From rostral to caudal levels, *Slc17a6* is expressed at high levels in the mitral and tufted cells of the main and accessory olfactory bulb, in the post olfactory eminence (an olfacto recipient region in the rostroventral telencephalon previously mistaken for part of the septum, see also (Endepols et al. 2005)), and in the lateral septum. Starting just posterior to the accessory olfactory bulb, high *Slc17a6* expression can be visualized as a longitudinal stripe of cells that extends to the most caudal levels of the telencephalic vesicle (Fig. 1D). Continuing posterior to the level of the interventricular foramen, *Slc17a6* is expressed in the lateral cellular prominence (a characteristic thickening of the cellular layer at the border of pallium and subpallium, that corresponds to the caudal part of the lateral amygdala) and along the entire ventral wall of the telencephalic vesicles, in regions previously described as parts of the urodele amygdala. In addition, low expression of *Slc17a6* is observed in the caudal region of the medial pallium and in the ventral region of the ventral pallium (VP), in line with the scRNAseq data. In comparison, *Gad1*, a marker of GABAergic neurons, is expressed rostrally in olfactory bulb interneurons and caudally throughout the subpallium (Fig. 1D). Interestingly, we observed that *Gad1*-expressing cells flank the *Sox6*-positive ventral pallium (Woych et al. 2022), at the same level of the highly-expressing *Slc17a6* neurons (Fig. 1E). In more posterior sections, *Slc17a6* and *Gad1* are clearly coexpressed in the lateral cellular prominence (Fig. 1D).

Together, we hypothesize that the lateral stripe of cells expressing high levels of *Slc17a6* corresponds to the lateral amygdala (LA) and includes both glutamatergic (pallial) and GABAergic (subpallial) neurons. We propose to refer to these glutamatergic neuron types as lateral pallial amygdala, to distinguish them from the GABAergic neurons they are intermingled with.

To investigate this further and identify the boundary between the ventral pallium, the lateral amygdala and the supallium, we analyzed the expression of *Gad1* (GABAergic), *Sox6* (ventral pallium), *Tac1* (striatum), *Penk* (striatum and TEGLU35) and *Lhx9* (TEGLU35) (Fig. 1F) (Woych et al. 2022). At mid-telencephalic levels, *in situ* hybridization on brain sections shows *Sox6* expression in the ventral pallium. Additionally, *Tac1* and *Penk* are both expressed at high levels in the striatum. However, *Penk* is also detected at lower levels in a band of cells just dorsal to the striatum (Fig. 1F). Similar to the *Slc17a6* stripe, this band of cells starts posterior to the accessory olfactory bulb and ends in the caudal telencephalon. We conclude that these weakly-expressing *Penk* cells correspond spatially to the stripe expressing high levels of *Slc17a6*. Confirming this, we find that *Penk* and *Slc17a6* are coexpressed in cluster TEGLU35, indicating the existence of *Penk*-expressing glutamatergic neurons that are not part of the GABAergic striatum (Fig. 1C). Supporting this conclusion further, *Lhx9*, a well-characterized marker of the lateral amygdala (Moreno et al. 2004; Garcia-Calero and Puelles 2021), is also expressed in TEGLU35 and stains neurons that occupy the same position as the weakly-expressing *Penk* cells (Fig. 1C,F).

The spatial distribution of cells expressing these genes, in combination with the scRNAseq data, also clarify the boundary between ventral pallium and pallial amygdala. Neither *Penk* nor *Lhx9* are coexpressed with *Sox6* in any cluster of the scRNAseq dataset. In contrast, some cells in the ventral pallium clusters (TEGLU19, 22-24) do co-express *Sox6* and *Slc17a6*, consistent with the HCR data, where the *Slc17a6* expression domain extends more dorsally than *Penk* and *Lhx9* expression (Fig. 1C,D). We conclude that cluster TEGLU35 corresponds to the lateral amygdala as defined by Moreno, Gonzales and colleagues, also referred to as the striato-pallial transition area by Marin and colleagues (Marín et al. 1997b, 1997a; Moreno and González 2004; Moreno et al. 2004; Laberge et al. 2006). Taken together, these data define the boundaries of the pallial amygdala at mid-telencephalic levels: rostrally, with the accessory olfactory bulb, dorsally, with the ventral pallium, and ventrally with the striatum.

### The *Pleurodeles* amygdala includes molecularly distinct types of glutamatergic neurons

To systematically explore the molecular diversity of the pallial amygdala, we subclustered glutamatergic amygdala cells (Fig. 2A). Some clusters were found to contain multiple distinct cell subtypes, and so we split clusters TEGLU25, TEGLU35, and TEGLU38 (indicated as TEGLU25.1 and 25.2, TEGLU35.1 and 35.2, and TEGLU38.1 and 38.2, respectively) (Fig. 2A-C). Together, our analysis of 1,417 single-cell transcriptomes suggests the existence of at least 10 distinct glutamatergic neuron types in the amygdala of *Pleurodeles*. To determine the exact spatial distribution of these amygdala neuron types, we drew on the scRNAseq dataset to identify cell type specific marker genes (Fig. 2B,C, see below and Fig. 1C).

**Fig. 2.**
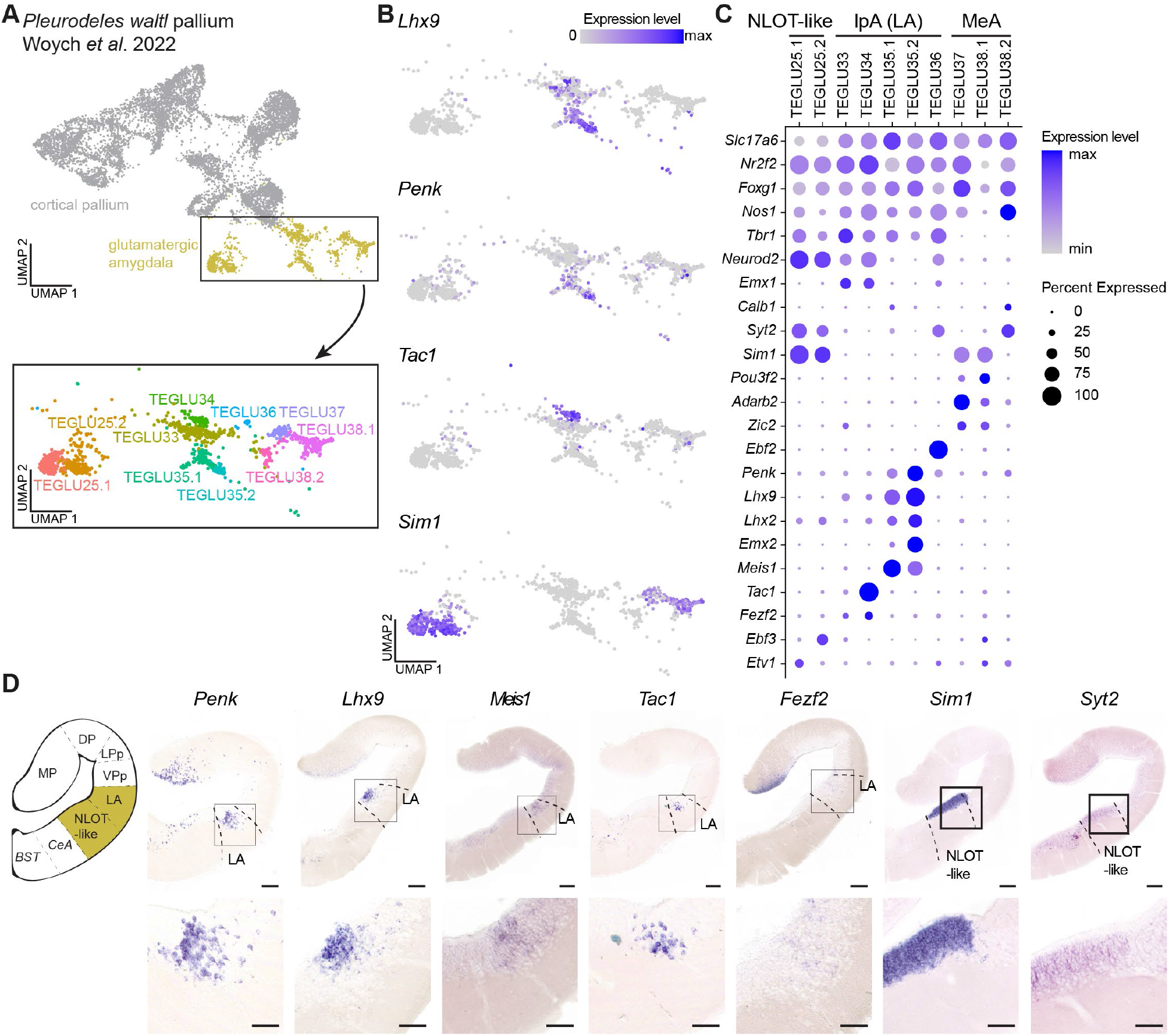
Neuronal heterogeneity in the *Pleurodeles* pallial amygdala. **(A)** UMAP plot of telencephalic glutamatergic neurons with cells from the amygdala highlighted in yellow. The magnified UMAP on the bottom shows the annotation of the different neuron types in the glutamatergic amygdala. **(B)** UMAP plots showing single cells colored according to the expression of *Lhx9, Penk, Tac1* and *Sim1*, illustrating the cellular diversity of the *Pleurodeles* amygdala. **(C)** Dot plot showing the expression of key marker genes defining distinct neuron types in the glutamatergic amygdala. **(D)** From left to right: schematic of a coronal section through the *Pleurodeles* telencephalon, expression of *Penk, Lhx9, Meis1, Tac1* and *Fezf2* in the lateral amygdala and of *Sim1* in the NLOT-like area. Boxes in upper panels indicate magnified regions in lower panels. Scale bars represent 200 um in upper panels and 100 um in lower panels. Abbreviations: BST, bed nucleus of the stria terminalis; CeA, central amygdala; DP, dorsal pallium; lpA, lateral pallial amygdala; LA, lateral amygdala; LPp, posterior lateral pallium; MP, medial pallium; NLOT-like, nucleus of the olfactory tract-like; TEGLU, telencephalic glutamatergic; VPp, posterior ventral pallium.

Focusing on the lateral pallial amygdala, our data show that *Lhx9* and *Meis1* are expressed at different levels in clusters TEGLU35.1 and 35.2 (Fig. 2B,C). *In situ* hybridization for these genes shows that they are expressed in the *Slc17a6*/*Penk* stripe of the lateral amygdala (Fig. 2D), recognizable in sections posterior to the intraventricular foramen as the lateral cellular prominence. The neuropeptide precursor gene *Tac1* is a specific marker of cluster TEGLU34. Expression analysis shows its presence in a subset of cells located in the lateral amygdala, further away from the ventricle compared to *Lhx9*. These *Tac1*-positive cells, or a subset of them, also express *Fezf2* (Fig. 2C,D). Additionally, *Calb1*, detected in cluster TEGLU35.1, is expressed in scattered cells of the lateral amygdala (Fig. 2C and (Morona and González 2008)).

Next, we investigated the expression of the transcription factor *Sim1*, expressed at high levels in clusters TEGLU25.1 and TEGLU25.2 and at low levels in clusters TEGLU37 and TEGLU38.1. To distinguish between these clusters, we also mapped the expression of *Syt2* (*Synaptotagmin 2*), a gene co-expressed with *Sim1* only in clusters TEGLU25.1 and TEGLU25.2 (Fig. 2C,D). *In situ* hybridization on brain sections and whole mount preparations show expression of *Sim1* at high levels in a large periventricular population of cells, along the ventrocaudal wall of the telencephalic vesicle, overlapping with the *Syt2* expression domain (Fig. 2D, Supplementary File 1). This population is immediately caudal to the striatum, medial to the *Lhx9*/*Penk* sector of the lateral amygdala, and dorsolateral to the central amygdala (see also Fig. 1D). A previous analysis of transcriptomic similarity across telencephalic clusters indicated that TEGLU25 is more similar to the ventral pallium than to other clusters from the pallial amygdala (Woych et al. 2022). However, the *Sim1* expression pattern clearly indicates that this cluster maps to a region of the amygdala, and not to the cortical pallium. In addition, unlike the lateral amygdala clusters, TEGLU25 is expressing a series of genes also detected in the mouse nucleus of the lateral olfactory tract (NLOT2), such as *Tbr1, Lhx2, Nr2f2* (*Coup-tf2*), *Etv1, Ebf3, NeuroD2*, and *NeuroD6* (Remedios et al. 2007; Tang et al. 2012; Garcia-Calero et al. 2020; Aerts and Seuntjens 2021). In light of this, here we will refer to this newly-defined amygdalar region as the NLOT-like area (see also Discussion).

Taken together, these results indicate that the lateral amygdala is heterogeneous, and includes multiple neuron types. We refer to the glutamatergic cells of clusters TEGLU33-TEGLU36 as the lateral pallial amygdala (lpA), in order to distinguish these cell types from the GABAergic cells of this region. The *Sim1* expression of clusters TEGLU25.1 and TEGLU25.2 is mapped to a region here called the NLOT-like area. While all the glutamatergic neuron types in the amygdala express *Slc17a6, Nr2f2, Foxg1* and *Nos1*, clusters TEGLU37-TEGLU38 are devoid of *Tbr1* expression (Fig 2B). As described below, these two clusters correspond to glutamatergic cells sampled from the medial amygdala (MeA).

### Expression of *Sim1* and *Otp* in the medial amygdala of *Pleurodeles*

From our analysis of the glutamatergic amygdala dataset, we found *Sim1* expression at low levels in clusters TEGLU37 and TEGLU38.1 (Fig. 2B). In these clusters, *Sim1* expression overlaps with high expression of the transcription factor *Otp* (Fig. 3A). Previous work in mammals (García-Moreno et al. 2010), lizards, and birds (Metwalli et al. 2022) identified a population of *Sim1+ Otp+* neurons that migrate from the periventricular hypothalamus to the medial amygdala; these neurons are born in a territory called telencephalon-opto-hypothalamic domain (TOH) (Morales et al. 2021) and migrate through the hypothalamo-amygdala corridor (HyA) (Garcia-Calero et al. 2021). We asked whether *Sim1+ Otp+* neurons also migrate from the periventricular hypothalamus to the medial amygdala in *Pleurodeles*. Therefore, analyzed the expression of *Otp* and *Sim1* using whole-mount *in situ* hybridization in the *Pleurodeles* brain (Fig. 3B,C, Supplementary File 2). As visualized in optical horizontal sections through the telencephalon, *Otp*-expressing cells lie in the same position as the weakly-expressing *Sim1* cells in the medial amygdala. Using 3D rendering, we observed that the *Otp* expression domain is a continuous stripe of cells that goes from the periventricular hypothalamus to the medial amygdala, and terminates just posterior to the cells expressing high levels of *Sim1* in the NLOT-like area described above (Fig. 3C, Supplementary File 3).

**Fig. 3.**
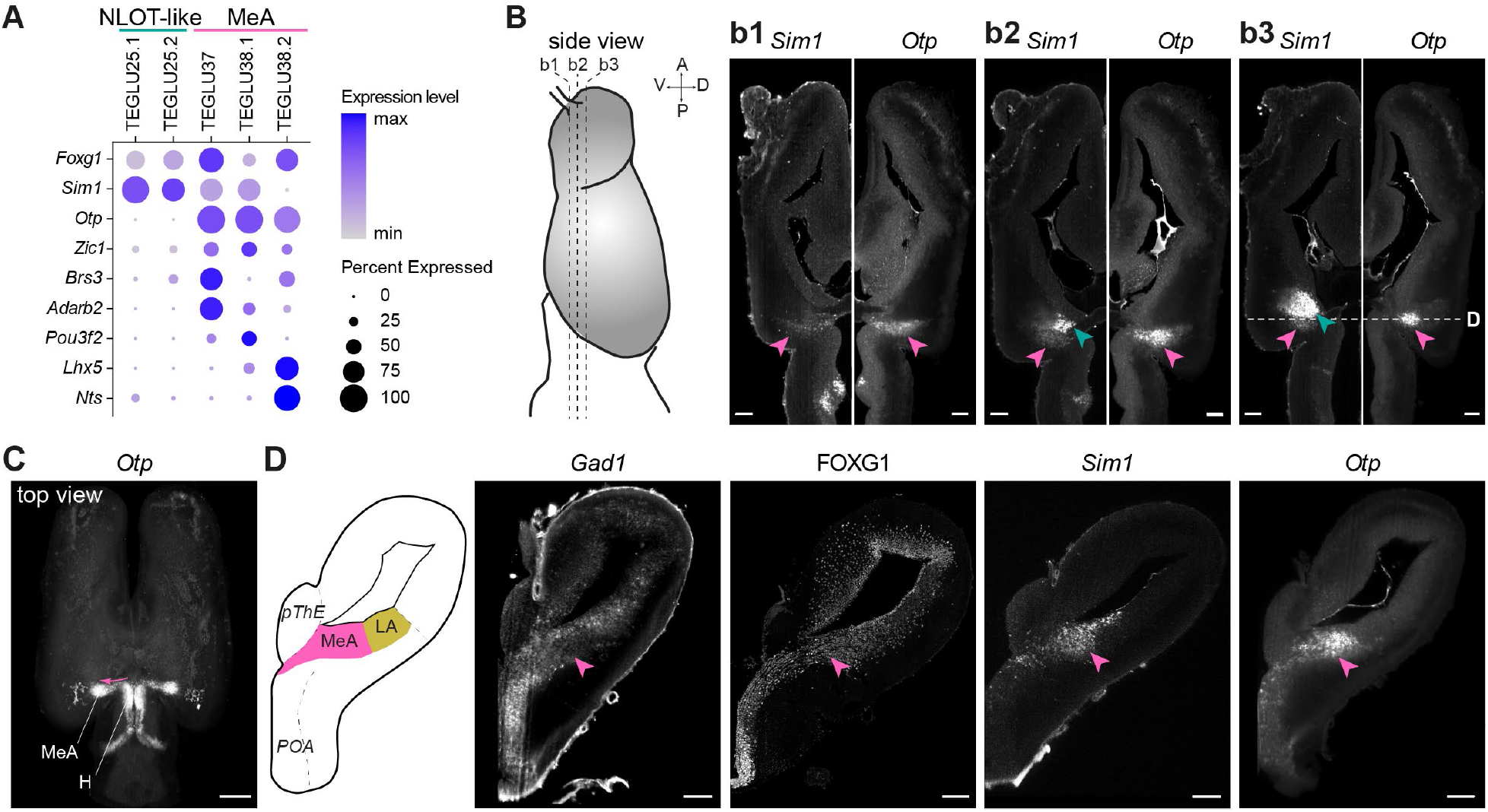
The hypothalamus-amygdalar corridor in *Pleurodeles*. **(A)** Dot plot showing differentially expressed genes in NLOT-like cells and the MeA. **(B)** Left: schematic of the side view of the salamander telencephalon (gray) with dashed lines indicating the optical sectioning plane for images on the right. Right: optical horizontal sections after whole-mount HCR ISH, clearing and light sheet imaging for *Sim1* and *Otp*. Pink and teal arrowheads indicate the medial amygdala and the NLOT-like area, respectively. Dashed line in b3 indicates the sectioning plane for D. Scale bars represent 200 um. **(C)** 3D rendering (Imaris viewer) of *Otp* expression in the *Pleurodeles* brain. Pink arrow indicates a continuum of *Otp+* cells from the periventricular hypothalamus to the medial amygdala. Scale bar represents 500 um. **(D)** From left to right: schematic of a coronal section at the level of the caudal telencephalon, expression of *Gad1*, presence of FOXG1, and expression of *Sim1* and *Otp*. Pink arrowheads indicate the medial amygdala. Scale bars represent 200 um. Abbreviations: LA, lateral amygdala; H, hypothalamus; MeA, medial amygdala; NLOT-like, nucleus of the olfactory tract-like; POA, preoptic area; pThe, prethalamic eminence; TEGLU, telencephalic glutamatergic.

According to our scRNAseq data, neurons in clusters TEGLU37 and TEGLU38 express *Zic1* and *Lhx5*, markers that are also found in the frog medial amygdala (Fig. 3A)(Jiménez and Moreno 2021). Furthermore, these neurons express *Foxg1* but not *Tbr1* (Fig. 2C), similarly to the *Sim1+* neurons that populate the mouse medial amygdala (Garcia-Calero et al. 2021). Finally, we find that the *Bombesin-like receptor 3* gene (*Brs3*), a specific marker of *Sim1+* neurons in the mouse medial amygdala, is also expressed in clusters TEGLU37 and TEGLU38.2 (Fig. 3A) (Xiao et al. 2020). Taken together, these data indicate the existence in *Pleurodeles* of *Sim1+ Otp+* neurons that are originating from the periventricular hypothalamus and populate the medial amygdala, and with a molecular profile similar to the *Sim1+ Otp+* neurons of the mouse medial amygdala.

## Discussion

### Organization of the amygdala in the urodele amphibian *Pleurodeles waltl*

In mammals, the amygdala is a heterogeneous collection of nuclei composed of neurons with diverse developmental origins and complex migratory histories. Reconstructing the evolution of the amygdala is further complicated by the fact that the amygdala is part of the caudal telencephalon, a brain region that underwent substantial changes during vertebrate evolution. At a first glance, amphibians may seem easy to analyze because they have a “simple” brain. However, the absence of clear neuroanatomical landmarks in the amphibian telencephalon, where distinct neuron types lay next to each other in a continuous periventricular cell layer, complicates the identification of boundaries and subregions. Here, we overcome this problem by using scRNAseq data as the starting point for the unbiased identification of amygdalar neuron types. The transcriptomics data confirm that amygdalar glutamatergic neurons are molecularly distinct from glutamatergic neurons of the cortical pallium (Woych et al. 2022). The differences arise mostly from the expression of genes in different combinations, rather than from the presence of a large number of region-specific markers, as it is the case for all subdivisions of the pallium. Furthermore, these data corroborate the distinction between a ventral pallium territory (labeled by the expression of the transcription factor *Sox6*) and the adjacent pallial amygdala.

Interestingly, glutamatergic neurons in the pallial amygdala and cortical pallium differ in their expression of vesicular glutamate transporters. In the amygdala, glutamatergic neurons express *Slc17a6* (*Vglut2*), whereas in the cortical pallium, glutamatergic neurons express *Slc17a7* (*Vglut1*). Exceptions to this rule include the weak expression of *Slc17a6* in parts of the ventral pallium, and the strong expression of *Slc17a7* in the newly-defined NLOT-like area (cluster TEGLU25). A similar dichotomy of vesicular glutamate transporter expression has been documented in mammals, where *Slc17a7* is expressed in the cerebral cortex (including the piriform cortex and the hippocampus) and in the cerebellum, while *Slc17a6* is expressed in subcortical regions and in the amygdala (Fremeau et al. 2001; Vigneault et al. 2015). Remarkably, in the turtle *Trachemys scripta, Slc17a6* and *Slc17a7* are coexpressed in all pallial glutamatergic neurons (Tosches et al. 2018), suggesting that the regulation of these two transporters in the pallium has changed during tetrapod evolution.

The expression of *Slc17a6* helps to define the boundaries of the amygdala in urodeles. Mapping the expression of *Slc17a6* and *Gad1* reveals a complex regional distribution of glutamatergic and GABAergic neurons. Our data suggest that glutamatergic and GABAergic neurons are intermingled in the lateral amygdala (named by Moreno and Gonzales (Moreno and González 2004; Moreno et al. 2004)), as clearly indicated by the expression of both *Slc17a6* and *Gad1* in the lateral cellular prominence. Furthermore, the sharp boundary of *Slc17a6* and *Gad1* expression in the ventrocaudal telencephalic wall revealed the presence of a new region, the NLOT-like area, nested between the striatum anteriorly and the medial amygdala posteriorly. Finally, we confirm that the medial amygdala, which occupies the most caudal part of the telencephalon, is also a mixture of glutamatergic and GABAergic neuron types.

### Neuronal diversity in the urodele lateral amygdala and evolutionary implications

A peculiarity of the amniote amygdala is the presence of multiple nuclei. In amphibians, only one pallial nucleus (the lateral amygdala) and two subpallial nuclei (the medial amygdala and the central amygdala) have been reported in the literature. From a comparative perspective, this has two alternative explanations: either the amphibian amygdala is simpler than the amygdala of amniotes (because it is more primitive or as a result of secondary simplification), or neuronal and regional diversity were overlooked in the amphibian amygdala for the absence of clear morphological landmarks. The analysis of neuron types, enabled by the scRNAseq approach, reveals that although the amygdala in urodeles is not as complex as in amniotes, there is still more cellular diversity than previously anticipated.

In the lateral amygdala alone, we identified at least 5 distinct clusters or subclusters of glutamatergic neurons (clusters TEGLU33, 34, 35.1, 35.2 and 36). We propose to call these neuron types lateral pallial amygdala, to distinguish them from the GABAergic neurons they are intermingled with. These neurons are labeled by the differential expression of several markers, including *Lhx9, Meis1, Calb1, Tac1*, and *Penk*, which were all detected at the pallial-subpallial boundary (at mid-telencephalic levels) and in the lateral cellular prominence (from levels caudal to the intraventricular foramen), confirming the lateral amygdala assignment of these clusters. Importantly, the cell type analysis reveals that *Lhx9*, a marker that has been used to identify the lateral amygdala in amphibians (Moreno et al. 2004), is expressed only in a subset of the lateral amygdala glutamatergic neurons. Interestingly, some of these markers appear to label cells with different distances from the ventricle (superficial or deep), suggesting that distinct amygdalar glutamatergic types might be generated sequentially during development, as it is the case for the rest of the amphibian pallium (Moreno and González 2017).

The recent radial model for the mammalian pallial amygdala posits that the pallial amygdala develops from five radial sectors (lateral, basal, anterior, posterior, and retroendopiriform), and that distinct amygdalar nuclei are formed by neurons generated sequentially within each of these sectors (Garcia-Calero and Puelles 2021). For example, the anterior basomedial amygdala (BMA) and the Anterior Cortical Amygdala (ACo) are generated within the same radial domain, where ACo neurons are born first, and BMA neurons are born later (Garcia-Calero et al. 2020). The exact number of radial domains in the pallial amygdala of non-mammals, including amphibians, remains undetermined. On the basis of our data, we can speculate that five distinct radial units did not exist in tetrapod ancestors, because we cannot find evidence of radial units of the pallial amygdala in *Pleurodeles*. It is conceivable that new specialized subdivisions of the pallial amygdala emerged in amniote or in mammalian ancestors together with the expansion of the rest of the telencephalon. Work on amygdala development in a variety of vertebrate species will be needed to reconstruct the evolution of these radial domains.

Our scRNAseq data on the adult salamander add a cell type perspective to the comparison of amygdala nuclei across species. On the basis of hodological data, the amphibian lateral amygdala has been compared to several amygdalar nuclei in amniotes. The amphibian lateral amygdala is extensively connected with other telencephalic areas (septum, bed nucleus of the stria terminalis, striatum), receives inputs from the thalamus and the parabrachial area, and projects to the ventromedial hypothalamus through the stria terminalis (Moreno and González 2004; Woych et al. 2022). In addition, the amphibian lateral amygdala receives inputs from both the main and the accessory olfactory bulb (Moreno and González 2004; Laberge and Roth 2005; Woych et al. 2022). In amniotes, olfactory inputs and other sensory inputs relayed through the thalamus reach distinct sets of amygdala nuclei. In mammals, the superficial cortical amygdala (early-born neurons according to the radial model) receives olfactory inputs, whereas the basolateral complex (lateral, basolateral and basomedial amygdala, late-born neurons in the radial model) receives multimodal thalamic inputs (lateral amygdala). The latter is reciprocally connected with the rest of the telencephalon, and projects to the ventromedial hypothalamus and other extra-telencephalic targets. In lizards, olfactory inputs reach the posterior lateral cortex and the nucleus sphericus, whereas the dorsolateral amygdala (DLA) and other parts of the posterior DVR have connections analogous to the mammalian basolateral complex (Voneida and Sligar 1979; Bruce and Neary 1995; Novejarque et al. 2004; Norimoto et al. 2020). These observations can be explained by at least two alternative hypotheses. Each neuron in the amphibian lateral amygdala might integrate both olfactory and non-olfactory information, and the processing of these modalities by distinct neurons and amygdalar nuclei may have evolved only in amniotes (Moreno and González 2007b). Alternatively, distinct neurons in the amphibian lateral amygdala would belong to separate subcircuits, which became segregated in distinct nuclei in amniotes. The surprising cellular heterogeneity of the *Pleurodeles* lateral amygdala supports the latter scenario. *Lhx9*, a transcription factor used previously to support the homology of the amphibian lateral amygdala with amniote pallial amygdala nuclei, is expressed only in a subset of the *Pleurodeles* lateral amygdala glutamatergic neurons. In E12.5 mice, *Lhx9* is expressed at high levels in the anterior radial unit, which includes BMA and ACo, but not in the basal and lateral units (precursors of the basolateral and lateral amygdalar nuclei, among others) (Abellán et al. 2014; Garcia-Calero and Puelles 2021). Interestingly, BMA and ACo not only develop from the same radial unit, but are also considered part of the same functional system (Petrovich et al. 1996): ACo integrates inputs from the main and the accessory olfactory bulb, and is reciprocally connected with the BMA (Cádiz-Moretti et al. 2017); the BMA instead is connected with olfactory cortical areas and sends projections to the ventromedial hypothalamus through the stria terminalis (Petrovich et al. 1996). In contrast, the mammalian basolateral amygdala (BLA), which develops from the basal unit (not to be confused with the mammalian lateral amygdala, a derivative of the lateral unit, (see (Garcia-Calero and Puelles 2021) for terminology) expresses a distinct set of transcription factors, including the specific markers *Fezf2* and *Etv1* (Hirata-Fukae and Hirata 2014; O’Leary et al. 2020). In *Fezf2* mutant mice, BLA glutamatergic neurons die after cell cycle exit, suggesting that *Fezf2* is a terminal selector transcription factor for BLA identity (Hirata-Fukae and Hirata 2014; O’Leary et al. 2020). In *Pleurodeles*, the lateral amygdala region includes *Fezf2*-expressing neurons (clusters TEGLU33 and 34), which are distinct from the *Lhx9*-expressing neurons described above. However, these neurons do not coexpress *Etv1*, according to our scRNAseq and *in situ* hybridization data. In reptiles, neurons co-expressing *Etv1* and *Fezf2* have been identified in the pDVR of turtles and lizards (Tosches et al. 2018; Norimoto et al. 2020), suggesting that the gene regulatory program that establishes the co-expression of these two transcription factors exists only in amniotes, but not in amphibians (or at least not in salamanders; see (Jiménez and Moreno 2021)). Additional cellular transcriptomics data in mammals, birds, reptiles, and other amphibians are needed to understand the evolutionary relationships of neuron types in the amphibian lateral amygdala, reptilian and avian pDVR, and mammalian cortical amygdala and basolateral complex.

Finally, the presence of GABAergic neurons in the lateral amygdala of *Pleurodeles* may provide new insights on the evolution of the mammalian intercalated cells (ITCs). In adult mice, ITCs surround the lateral and the basolateral amygdala, and populate the intercalated mass of the amygdala (IA), located between pallial and the subpallial amygdalar nuclei. Fate mapping experiments showed that unlike other amygdala interneurons, which originate in the ventral lateral ganglionic eminence (LGE), ITCs develop from the dorsal LGE (dLGE), and therefore share their developmental origin with medium spiny neurons of the striatum (Waclaw et al. 2010; Bandler et al. 2022). During development, the mammalian dLGE is immediately ventral to the pallium, in a position that is topologically homologous to the most dorsal part of the subpallium in amphibians. In mammals, ITCs migrate after birth in a ventrocaudal direction, through the lateral migratory stream, to reach their final destination in the amygdala. Mouse ITCs express a unique set of transcription factors and marker genes, and the analysis of these markers in our scRNAseq data suggests the existence of ITC-like neurons in *Pleurodeles*. In mice, ITCs are *Penk*-positive, but unlike striatal *Penk* cells, they express the transcription factor *Foxp2* instead of its paralog *Foxp1*. Furthermore, ITCs are distinct from the striatum for the expression of the transcription factors *Tshz1* and *Pax6* (*Kuerbitz et al. 2018*); they also express *Sp8* and *Meis2* (like the striatum), but lack the expression of the striatal marker *Isl1*. In turtles and lizards, LGE-derived neurons with a ITC-like expression profile have been identified; these neurons co-express *Foxp2, Penk, Tshz1, Pbx3*, and *Meis2*, and the *Foxp2* expression pattern indicates their location in the pDVR (Tosches et al. 2018). In *Pleurodeles*, a subcluster of *Penk*-expressing GABAergic neurons (TEGABA55, (Woych et al. 2022)) coexpresses *Penk, Foxp2, Tshz1, Pax6, Pbx3*, and *Meis2*, but is negative for *Foxp1* and *Isl1* (Supplementary Fig. S1). Both *Penk* and *Pax6* are expressed in the lateral amygdala (Joven et al. 2013), suggesting that the GABAergic neurons of the lateral amygdala correspond to this subcluster. Taken together, these data suggest that ITC GABAergic neurons originating from the dLGE are ancestral in tetrapods.

### *Sim1*-expressing neuron types in the urodele amygdala and evolutionary implications

Here we describe a new division of the amygdala in *Pleurodeles*, which we provisionally name NLOT-like area in light of its gene expression profile (cluster TEGLU25). The scRNAseq analysis indicates that these neurons are extremely different, at a molecular level, from neurons in the lateral amygdala. The most distinctive feature of cluster TEGLU25 is the expression of some cortical pallium markers, such as *Slc17a7, NeuroD2*, and *NeuroD6*, which initially misled us to think that TEGLU25 was part of the cortical pallium too. However, the expression pattern of the transcription factor *Sim1* in the adult brain shows unambiguously that TEGLU25 neurons are part of the amygdala, with a lateral boundary with the lateral amygdala, a rostral boundary with the striatum and a caudal boundary with the medial amygdala. This region of the amphibian brain is considered part of the medial amygdala in the literature (Moreno and González 2007c). Our scRNAseq data suggest an alternative interpretation. The expression of several genes reveal similarities of these *Pleurodeles Sim1*-expressing cells with cells in the mammalian nucleus of the lateral olfactory tract (NLOT). Like the mammalian NLOT, this region of the urodele telencephalon is receiving inputs from the main olfactory bulb (Laberge and Roth 2005; Moreno and González 2007c). In mice, *Sim1* is expressed in layer 2 of the NLOT (NLOT2), together with other markers such as *Tbr1, Lhx2, Etv1, NeuroD2, NeuroD6, Zic2*, and *Dach1* (Remedios et al. 2007; Aerts and Seuntjens 2021; Garcia-Calero et al. 2021). Mouse NLOT2 neurons arise from multiple progenitor domains: the hypothalamus (source of *Sim1*+ neurons) and the posterior amygdalar radial unit (source of TBR1+ neurons) (Remedios et al. 2007; Garcia-Calero et al. 2020). From these birthplaces, *Sim1*+ (mRNA) and TBR1+ (protein) neurons are thought to migrate rostrally along converging routes to populate NLOT2 (Remedios et al. 2007; Aerts and Seuntjens 2021; Garcia-Calero et al. 2021). Interestingly, the *Pleurodeles* TEGLU25 cluster co-expresses *Sim1* and *Tbr1* mRNAs, together with other markers of NLOT2 (in TEGLU25.1, TEGLU25.2 or both) such as *Lhx2, Nr2f2* (*Coup-tf2*), *Etv1, Ebf3, NeuroD2*, and *NeuroD6* (but not *Zic2* and *Dach1*) (Remedios et al. 2007; Tang et al. 2012; Garcia-Calero et al. 2020; Aerts and Seuntjens 2021). *Sim1*+ neurons have been found also in the chick LOT nucleus, putative homolog of the mammalian NLOT ((García-Moreno et al. 2010; Garcia-Calero et al. 2021; Morales et al. 2021; Metwalli et al. 2022)). These data raise questions on the developmental origins of NLOT-like neurons in *Pleurodeles*. The expression of “cortical pallium” markers (*Tbr1, NeuroD2, NeuroD6*) may suggest a pallial origin; however, the expression of *Sim1* is in conflict with this interpretation. Furthermore, some “cortical pallium” markers are expressed in the mouse preoptic area and hypothalamus too (*Tbr1*: (Bulfone et al. 1995; Puelles et al. 2000; Romanov et al. 2020), *NeuroD6*: (Caqueret et al. 2006)), although it cannot be excluded that this is the result of cell migration from the pallium or the prethalamic eminence, at least in some cases. Further analysis of developmental stages in *Pleurodeles* and other amphibians is needed to determine the developmental origins of NLOT-like cells.

The NLOT-like area is distinct from a second population of *Sim1*-expressing neurons immediately caudal to it. These neurons co-express *Sim1* and *Otp* (clusters TEGLU37 and TEGLU38) and lack *Tbr1* expression; *Otp* whole-mount *in situ* hybridization reveals a continuous population of neurons from the periventricular hypothalamus to the caudal telencephalon. For their position and their characteristic molecular profile, we propose that these neurons are homologous of the *Sim1+ Otp+* neurons of the mammalian medial amygdala, recently also identified in sauropsids (García-Moreno et al. 2010; Garcia-Calero et al. 2021; Morales et al. 2021; Metwalli et al. 2022). Unlike in the amniote situation, where distinct *Sim1+ Foxg1+* and *Sim1+ Foxg1*-neurons have been identified, our staining data indicate so far that the medial amygdala *Sim1+ Otp+* cells co-express *Foxg1* in *Pleurodeles*, suggesting that the amniote amygdala harbors a larger variety of neuron types. Taken together, our results establish that *Sim1+ Otp+ Foxg1+* neurons of the hypothalamic-amygdalar corridor (Garcia-Calero et al. 2021), also referred to as telencephalon-opto-hypothalamic (TOH) domain (Morales et al. 2021), were present in the last common ancestor of tetrapods and contributed glutamatergic neurons to the medial amygdala. The existence of *Otp+* neurons in the amygdala of zebrafish (Porter and Mueller 2020) suggests that this neuron type may trace back to vertebrate ancestors, although additional work in fish and cyclostomes is needed to test this hypothesis.

### The amygdala as an example of mosaic evolution in the brain

There are mixed opinions in the literature on the relevance of salamanders for brain evolution studies. Herrick (Herrick 1948) assumed that urodele amphibians retained the ancestral characters of the brains of tetrapod ancestors, while Roth et al (Roth et al. 1993) pointed out that the morphological simplicity of the salamander brain evolved secondarily as a result of paedomorphosis (retention of embryonic or juvenile characters in adulthood). These analyses, however, were limited to neuroanatomical traits, such as the migration of newly-born neurons away from the ventricle. Our molecular analysis of neuron types in *Pleurodeles* indicates that the salamander telencephalon hosts an unanticipated variety of neuron types, which can be readily compared to neuron types in amniotes (Woych et al. 2022). This suggests that brain morphology and neuronal diversity are partially uncoupled in evolution, and that brain structures with higher or lower degrees of morphological complexity can harbor homologous neuron types. The independent evolution of brain morphology and neuron types is an example of mosaic evolution, where biological traits evolve at different rates under different selective pressures (Northcutt and Kicliter 1980). In light of this, the simple organization of the salamander brain can be exploited to accelerate the discovery of neuron types and neural circuits shared across tetrapods, provided that their phylogenetic continuity can be demonstrated. In the case of the amygdala, we conclude that the amygdala of stem tetrapods had a heterogeneous repertoire of glutamatergic neurons, including distinct *Lhx9*-positive and *Lhx9*-negative neurons in the pallial amygdala, *Sim1*+ neurons putatively homolog to the NLOT, and conserved *Sim1+ Otp+* neurons in the medial amygdala. Further characterization of amygdalar neuron types in developmental and adult stages of amniotes and anamniotes is needed to reconstruct with higher precision the evolution of this incredibly complex part of the brain.

## Acknowledgements

The authors are grateful to J. Barber, S. Cook, and the Columbia University Institute of Comparative Medicine for animal care, E. Gumnit for help with cloning, and A. Ortega Gurrola for critical feedback on the manuscript. Light-sheet imaging was performed with support from L. Hammond and the Zuckerman Institute’s Cellular Imaging platform (NIH 1S10OD023587-01).

## Statement of Ethics

This study protocol was reviewed and approved by the Columbia University Institutional Animal Care and Use Committee (IACUC), approval number AC-AABF2564.

## Author Contributions

scRNAseq analysis: AD; anatomy and histology: JW, AD, ECJ; manuscript writing: MAT, AD, JW, with edits from ECJ; project management and supervision: MAT.

## Data Availability Statement

The scRNA sequencing data used in this study have been deposited in the Gene Expression Omnibus (GEO), with accession numbers GSE197701, GSE197722, GSE197796, GSE197807 and GSE198363, as reported in (Woych et al. 2022). The adult pallium dataset can be browsed on https://toscheslab.shinyapps.io/salamander_telencephalon/.

## Figures

**Supplementary File 1. *Sim1* expression in the entire *Pleurodeles* brain in optical coronal sections**.

Light-sheet imaging of an adult *Pleurodeles* brain, stained by HCR in situ hybridization with a *Sim1* probe, and cleared with iDisco.

**Supplementary File 2. *Otp* expression in the entire *Pleurodeles* brain in optical coronal sections**.

Light-sheet imaging of an adult *Pleurodeles* brain, stained by HCR in situ hybridization with an *Otp* probe, and cleared with iDisco.

**Supplementary File 3. 3D rendering of *Otp* expression in the *Pleurodeles* brain**. 3D rendering of the light-sheet data from Supplementary File 2, showing the continuum of *Otp* cells from the periventricular hypothalamus to the medial amygdala.

**Supplementary Fig. 1.**
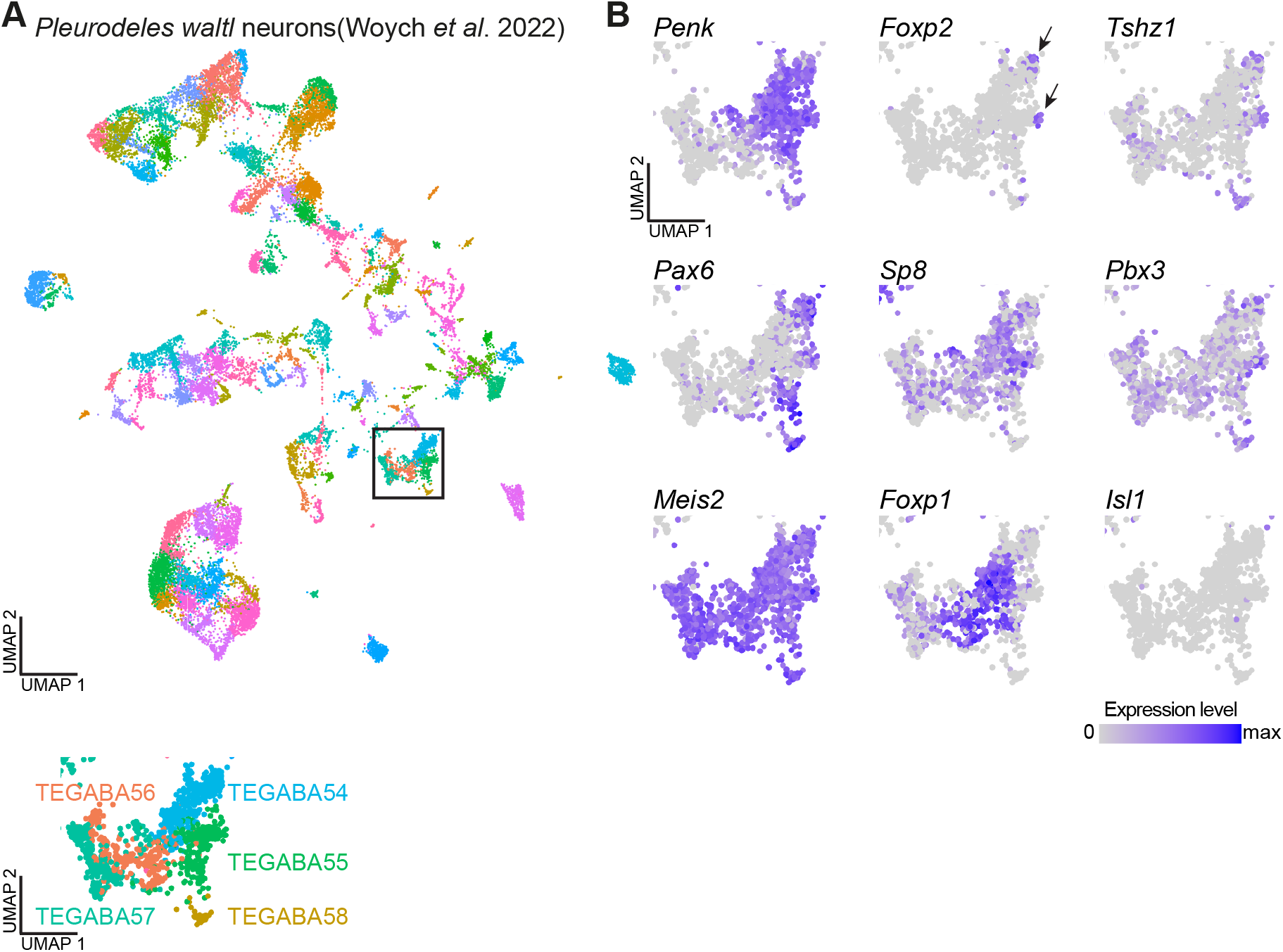
ITC-like cells in the *Pleurodeles* amygdala. **(A)** UMAP plot of the neuronal dataset, colored by cluster. Boxed area indicates magnified clusters on the bottom, highlighting TEGABA54-58. **(B)** UMAP plots showing single cells of clusters TEGABA54-58, colored according to the expression of *Penk, Foxp2, Tshz1, Pax6, Sp8, Pbx3, Meis2, Foxp1* and *Isl1*, illustrating *Pleurodeles* neurons with an ITC-like expression profile (arrows). Abbreviations: ITC, intercalated cells.

